# Dissociable Oscillatory Alpha and Aperiodic Neural Dynamics Underlie Atypical Attentional Control in Children With Attention-Deficit/Hyperactivity Disorder (ADHD)

**DOI:** 10.64898/2026.11.01.763095

**Authors:** Yuanjun Kong, Runhao Lu, Sung-Mu Lee, Manqi Zhou, Ziang Yu, Yung-Jui Lu, Xiaojun Sun, Ole Jensen, Li Sun, Yan Song

**Affiliations:** Key Laboratory of Adolescent Cyberpsychology and Behavior (CCNU), Ministry of Education, Wuhan 430079, China; Key Laboratory of Human Development and Mental Health of Hubei Province, School of Psychology, Central China Normal University, Wuhan 430079, China; Montreal Neurological Institute, Department of Neurology and Neurosurgery, McGill University, Montreal, Quebec H3A 2B4, Canada; Jefferson Moss Rehabilitation Research Institute, Thomas Jefferson University, Philadelphia 19027, USA; State Key Laboratory of Cognitive Neuroscience and Learning & IDG/McGovern Institute for Brain Research, School of Psychology, Beijing Normal University, Beijing 100875, China; Department of Experimental Psychology, University of Oxford, Oxford OX2 6GG, United Kingdom; Oxford Centre for Human Brain Activity, Wellcome Centre for Integrative Neuroimaging, University of Oxford, Oxford OX3 7JX, United Kingdom; Peking University Sixth Hospital & Peking University Institute of Mental Health, Beijing 100191, China; NHC Key Laboratory of Mental Health (Peking University) & National Clinical Research Center for Mental Disorders (Peking University Sixth Hospital), Beijing 100191, China; Beijing Key Laboratory of Consciousness Brain Computer Interface and Neuromodulation, Beijing Normal University, Beijing 100875, China

**Keywords:** ADHD, alpha oscillations, aperiodic activity, attentional control, distractor processing

## Abstract

Children with attention-deficit/hyperactivity disorder (ADHD) often have difficulty maintaining goal-directed attention when task-relevant and task-irrelevant information compete for processing resources. The present study investigated whether this difficulty reflects distinct abnormalities in demand-related attentional regulation and distractor-related neural responses. We recorded electroencephalography (EEG) signals from 45 children with ADHD and 45 typically developing (TD) children while they performed a visual detection task under low and high visual demands, with simultaneous task-irrelevant standard and deviant sounds. EEG power spectra were parameterized to separate oscillatory alpha activity from aperiodic activity, as putative indices of attentional regulation and cortical excitation/inhibition (E/I) balance, respectively. TD children showed adaptive demand-related alpha modulation, whereas children with ADHD showed an atypical alpha response pattern, including reduced frontal modulation and atypical posterior alpha enhancement; this demand-related group difference was observed in oscillatory alpha activity but not in aperiodic activity. In contrast, ADHD-related abnormalities in responses to auditory distractors were more clearly reflected in the aperiodic slope. Children with ADHD showed atypical distractor-related aperiodic modulation, suggesting reduced stability of neural background activity, potentially reflecting disrupted cortical E/I balance when irrelevant sounds had to be ignored. Furthermore, multivariate machine learning classification demonstrated that relying on these sensitive neural signatures yielded robust discrimination between children with ADHD and TD controls, with significant predictive performance emerging shortly after stimulus onset and sustaining throughout information processing. These findings suggest that oscillatory alpha activity and aperiodic neural dynamics show differential sensitivity to distinct attentional abnormalities in ADHD, with alpha oscillations primarily reflecting demand-related regulatory deficits and aperiodic activity primarily reflecting distractor-related neural instability.

## Introduction

Attention-deficit/hyperactivity disorder (ADHD) is a prevalent neurodevelopmental disorder characterized by symptoms including inattention, hyperactivity, and impulsivity, affecting approximately 7.2% of children worldwide [1]. Beyond these core behavioral symptoms, ADHD is frequently associated with impairments in the dynamic regulation of attentional resources, particularly when maintaining focus on task-relevant information under increasing task demands while simultaneously filtering out irrelevant distractors [2, 3]. Such situations require the flexible adjustment of attentional resources to match varying cognitive demand, as well as resilient responses to distracting environmental events [4–6]. Understanding the neural mechanisms underlying this demand-related modulation and distraction processing is therefore essential for clarifying the physiological basis of attentional dysfunction in children with ADHD.

Neural activity in the alpha band (8–12 Hz) has long been linked to the top-down regulation of visual attention [7–9]. According to the gating-by-inhibition framework [7], alpha oscillations shape the brain’s functional architecture by selectively silencing task-irrelevant pathways via power increases (event-related synchronization, ERS), while prioritizing goal-directed information flow through power decreases (event-related desynchronization, ERD) [10, 11]. As task difficulty increases, this gating mechanism becomes more pronounced, with alpha ERD becoming stronger in task-relevant regions, reflecting higher recruitment of attentional resources [12]. This gating mechanism is particularly evident in cross-modal tasks, where alpha ERS appears in brain areas processing irrelevant noise, while ERD appears in areas processing the attended information [13, 14]. For instance, Foxe et al. [15] demonstrated that alpha power over the occipital cortex increases when attention is directed toward the auditory component of an audiovisual stimulus, suggesting functional suppression of the visual system. Building on this, Fu et al. [16] found that anticipatory parieto-occipital alpha ERS effectively primes the brain for the subsequent deployment of attention to cued auditory stimuli.

In the context of ADHD, however, this flexible alpha-mediated control process appears to be significantly altered. Systematic reviews indicate that ADHD is characterized by an inability to adaptively modulate alpha activity in response to task-related cues or shifting goals [17–19]. Specifically, research shows that individuals with ADHD exhibit both attenuated alpha ERS and ERD across various cognitively demanding contexts, including visuospatial attention and working memory [20–22]. This suggests an underlying deficit in the dynamic allocation of neural resources needed to shield task-relevant processing [17, 23, 24], highlighting that alpha dynamics are sensitive to differences in demand-related attentional regulation in ADHD.

Beyond alpha oscillations, recent evidence increasingly suggests the important roles of aperiodic (1/f) activity in supporting attention and cognitive control [25–28]. Functionally distinct from rhythmic oscillations, aperiodic activity is theorized to reflect the global balance between neural excitation and inhibition (E/I balance) and the underlying level of neural excitability [29–33]. Critically, because traditional spectral analyses often conflate oscillatory and aperiodic signals, previous findings regarding alpha power shifts may be confounded by changes in the aperiodic background. Emerging research suggests that the aperiodic slope is not merely a static baseline but exhibits systematic, time-locked modulations following stimulus onset [31, 34], wherein shifts toward flatter slopes reflect increased excitation relative to inhibition, which likely track reactive neural responses to environmental events [35]. Specifically, the ability to adaptively modulate the cortical E/I balance may determine how effectively the brain maintains stability when confronted with unexpected distractors. However, while aperiodic abnormalities have been reported in several resting-state datasets in ADHD, research investigating task-related aperiodic dynamics remains exceedingly scarce, and the few existing studies have yielded discordant results (see Donoghue [36] for a review). Crucially, it remains largely unknown whether atypical aperiodic dynamics in ADHD are preferentially associated with an inability to regulate these reactive E/I shifts during environmental distraction. Given that ADHD is increasingly conceptualized as a context-dependent disorder of self-regulation [37], it is possible that sudden distractors under high cognitive demand might trigger a reactive destabilization of the cortical E/I balance, manifesting as transient fluctuations in the aperiodic slope.

Beyond clarifying these neural mechanisms, identifying objective physiological markers is of great clinical importance, as ADHD diagnosis remains primarily based on clinical interviews and symptom ratings, which may be influenced by subjective reports and clinical judgment [38, 39]. Accordingly, multivariate machine learning classification has been increasingly used to evaluate whether objective neurophysiological signatures can support ADHD–TD differentiation at the individual level [40]. Although integrating multiple features is often considered advantageous in machine-learning-based classification, greater classification accuracy does not necessarily come from including more features, because irrelevant or redundant features may compromise classification performance [41]. This highlights the need to identify neural features that are specifically sensitive to the cognitive processes most affected in ADHD. Thus, separating oscillatory alpha and aperiodic neural dynamics across distinct attentional processes may help identify more clinically informative markers for ADHD-TD classification.

In the present study, we utilized a cross-modal attention task that varied visual task demand while presenting task-irrelevant auditory distractors, thereby isolating demand-related regulation from distraction effects (Fig. 1A). By disentangling periodic and aperiodic components via a spectral parameterization algorithm [26], we sought to clarify whether these neural dynamics show distinct sensitivity to demand-related regulation and distraction effects in children with ADHD. Our hypotheses are as follows: First, we predicted that deficits in goal-directed demand modulation in ADHD would be reflected in oscillatory alpha activity, such that children with ADHD would exhibit atypical demand-related alpha modulation relative to TD controls as task demand increases. Second, we hypothesized that atypical processing of auditory distraction in ADHD would be reflected in aperiodic slope, with the ADHD group showing atypical cortical E/I balance in response to task-irrelevant sound changes. Finally, we reasoned that separating these neural markers would facilitate machine-learning-based classification with potential clinical relevance, hypothesizing that a model based on these theoretically sensitive neural features could distinguish children with ADHD from TD controls more effectively than models based on less sensitive features or feature combinations.

**Fig. 1.**
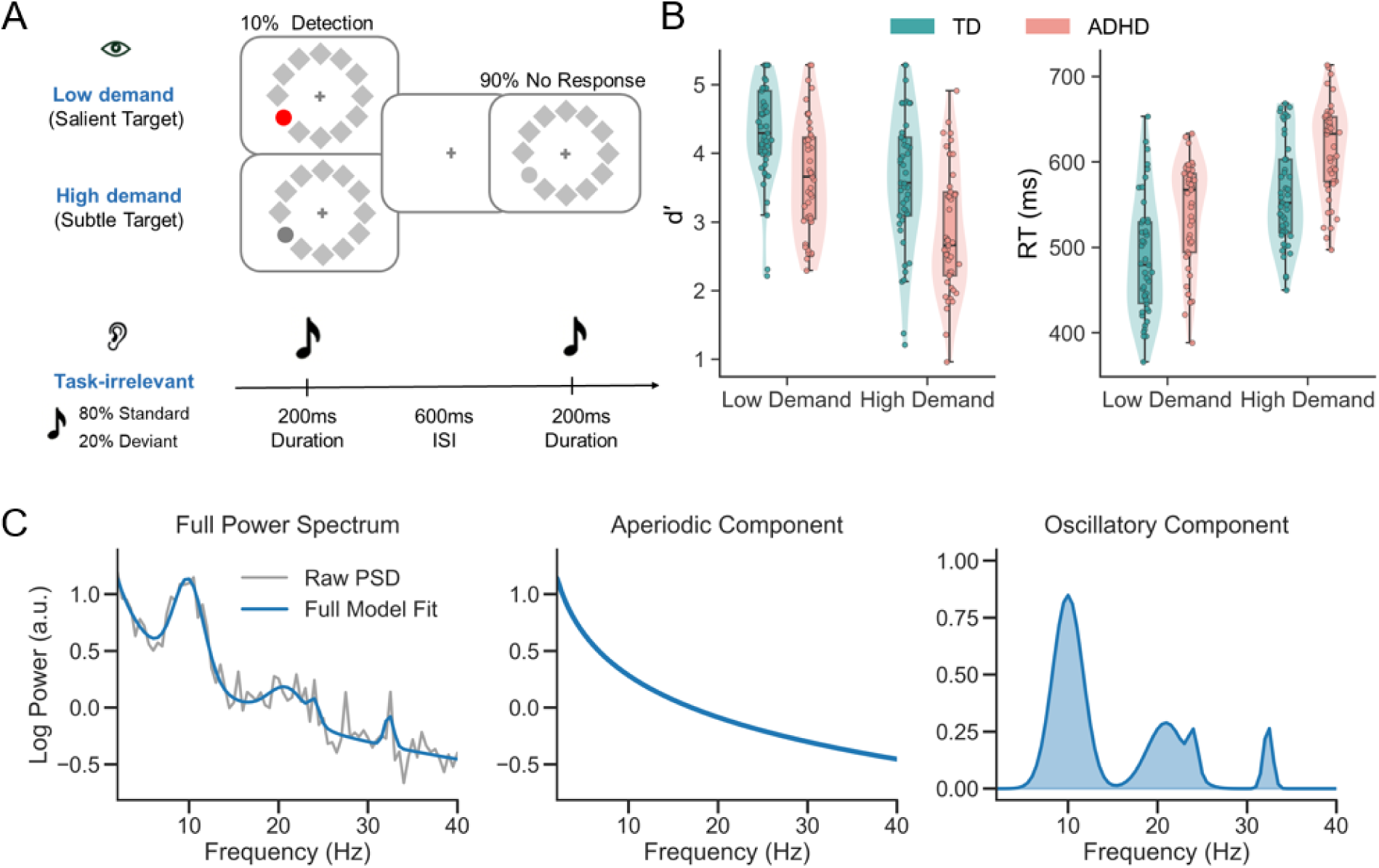
Experimental paradigm, behavioural performance, and decomposition of oscillatory and aperiodic activity. (A) Task paradigm. Participants performed a visual detection task under two demand conditions (Low demand: salient target; High demand: subtle target) while ignoring concurrent task-irrelevant auditory stimuli (80% standard and 20% deviant tones). Visual arrays and synchronized auditory stimuli were presented for 200 ms, followed by a 600 ms interstimulus interval (ISI). (B) Behavioural performance. Target discriminability (d′; left) and reaction time (RT; right) for the TD and ADHD groups under low and high visual task demand. Violin plots and boxplots illustrate the distribution and median/interquartile range of individual data points. (C) Parameterization of EEG power spectra. Representative illustration of the FOOOF decomposition used to dissociate oscillatory and aperiodic components from the raw power spectral density (PSD). Left: The full power spectrum (grey) and its corresponding full model fit (blue). Middle: The isolated aperiodic component reflecting the broadband 1/f background dynamics. Right: The isolated oscillatory components.

## Results

### Behavioural results

Participants performed a visual detection task under low and high attention demands (Fig. 1A), while task-irrelevant auditory standard and deviant tones were presented concurrently. Participants responded to infrequent visual targets and ignored the auditory stimuli.

To characterize behavioural performance in terms of target discriminability an d response speed, we analysed d′ and reaction time (RT) using mixed ANOVAs with group (ADHD vs. TD) and visual demand (low vs. high). For d′, there were significant main effects of group (*F*(1, 88) = 16.53, *p* < 0.001, *η*_p_^2^ = 0.158) and demand (*F*(1, 88) = 120.18, *p* < 0.001, *η*_p_^2^ = 0.577), with no group × demand interaction (*F*(1, 88) = 0.90, *p* = 0.344). TD children showed higher d′ than children with ADHD under both low and high demand. A similar pattern was observed for RT. There were significant main effects of group (*F*(1, 88) = 17.04, *p* < 0.001, *η*_p_^2^ = 0.162) and demand (*F*(1, 88) = 353.98, *p* < 0.001, *η*_p_^2^ = 0.801), but no group × demand interaction (*F*(1, 88) = 0.01, *p* = 0.934). Children with ADHD responded more slowly than TD children under both low and high demand. Together, these results indicate that the behavioural impairment in ADHD was evident in both reduced target discriminability and slower responses.

### Impaired demand-related oscillatory alpha modulation in ADHD

To elucidate the neural basis of potential deficits in demand-related attentional modulation in ADHD, we first examined how varying visual task demands modulated pure oscillatory alpha power (8–12 Hz) and the broadband aperiodic slope (Fig. 1C). Before spectral analysis, condition-specific ERPs were subtracted from each trial to minimize contributions from phase-locked evoked responses to the estimated oscillatory and aperiodic dynamics (see Methods). We evaluated this demand-related modulation by subtracting the neural metrics of the low-demand condition from those of the high-demand condition.

For oscillatory alpha power, we first analysed the time courses within our a priori spatial ROIs (Fig. 2A). These regions were selected to represent the functional specialization of the attentional network, with the frontal ROI serving as a proxy for top-down cognitive control and the posterior ROI reflecting the site of visual sensory modulation. In the frontal ROI, TD children exhibited a robust demand-related frontal alpha ERD, which emerged shortly after stimulus onset, reflecting the successful engagement of top-down cognitive control to facilitate task-relevant processing. Conversely, children with ADHD failed to exhibit this typical frontal ERD. Direct group comparison confirmed a significant divergence between the two groups in the frontal ROI from 188 to 540 ms (cluster-based permutation test: *p* < 0.05, two-tailed), potentially indicating an atypical demand-related modulation to task demand. Furthermore, we found that the atypical synchronization was particularly pronounced and sustained over the posterior region. This posterior ERS in children with ADHD under high cognitive demand (32 to 600 ms; cluster-based permutation test: *p* < 0.05, two-tailed) suggests potentially insufficient recruitment of visual sensory resources to support task performance under increased cognitive demand. To evaluate the spatial extent of these effects, we conducted exploratory whole-scalp cluster-based permutation testing on data averaged over the a priori 200–400 ms time window, a period typically associated with the active deployment of top-down attention and sensory gating [42]. Specifically, within-group topographical analyses confirmed that the demand-related alpha ERD in TD children was predominantly localized to a significant cluster over the frontal scalp. In contrast, the alpha ERS in children with ADHD was primarily clustered over posterior regions. Consequently, the direct group comparison revealed a widespread difference across the scalp (Fig. 2B).

**Fig. 2.**
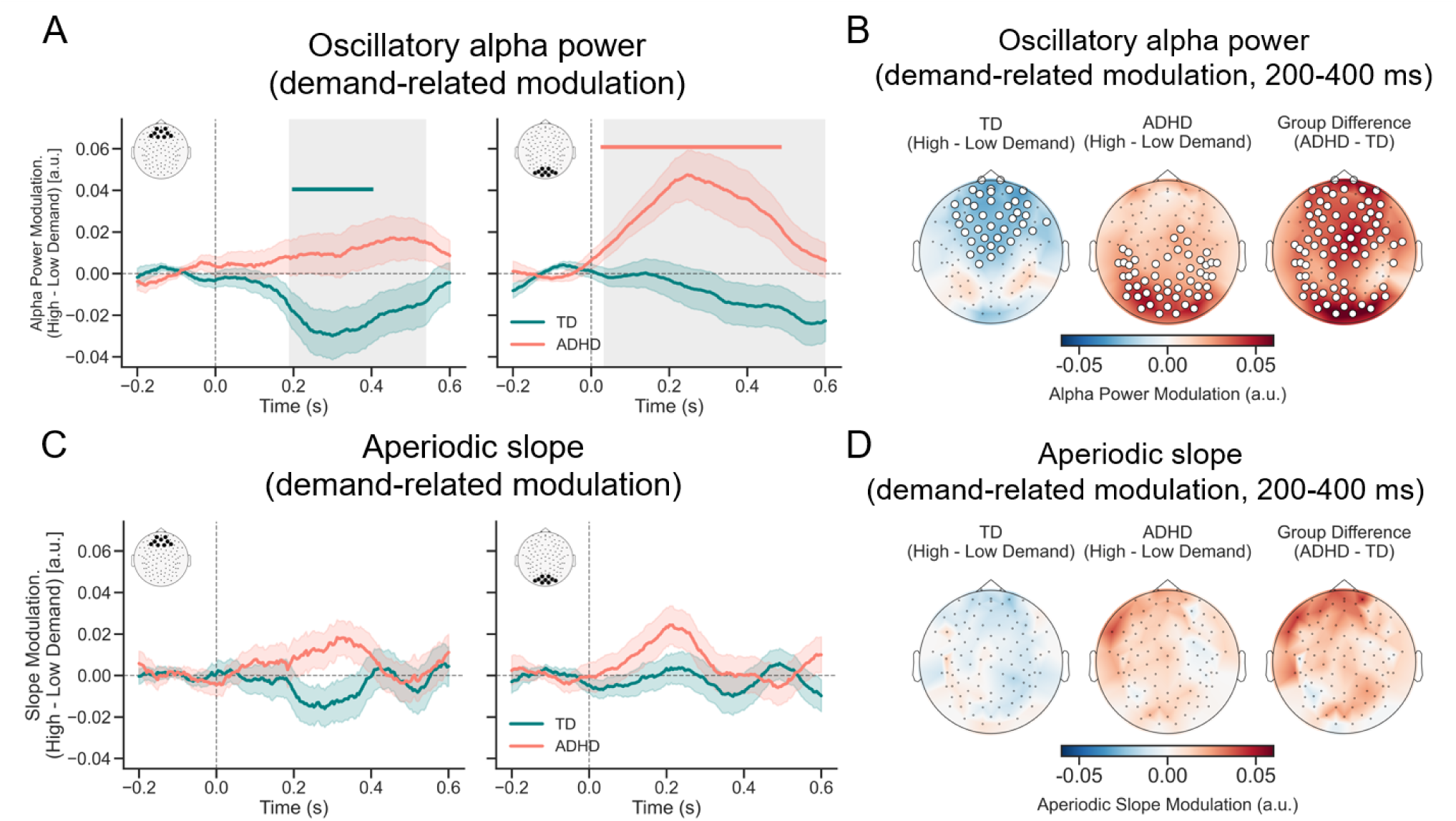
Impaired demand-related oscillatory alpha modulation in children with ADHD and its behavioural relevance. (A) Alpha power modulation to visual task demand (Time courses). Time courses of demand-related oscillatory alpha power modulation (High minus Low visual demand) within the *a priori* frontal (left) and posterior (right) regions of interest (ROIs). The inset topographical maps in the upper left corners illustrate the spatial locations of these predefined ROIs. Shaded error bands around the mean waveforms represent the standard error (SE). Solid coloured horizontal lines indicate significant within-group temporal clusters differing from zero. Grey shaded areas denote significant time windows for the group interaction (ADHD vs. TD) identified via cluster-based permutation testing. (B) Alpha power modulation to visual task demand (Topographical maps). Spatial distribution of alpha power modulation for each group and the statistical group difference (ADHD minus TD) averaged over the 200–400 ms time window. White dots highlight electrodes belonging to significant spatial clusters. (C) Aperiodic slope modulation to visual task demand (Time courses). Time courses of demand-related aperiodic slope modulation (High minus Low visual demand) within the frontal and posterior ROIs. (D) Aperiodic slope modulation to visual task demand (Topographical maps). Spatial distribution of demand-related aperiodic slope modulation averaged over the 200–400 ms time window.

Crucially, this atypical demand-related modulation in the ADHD group was specific to periodic alpha oscillations. When we examined the aperiodic slope, we found no significant demand-related group differences. Time-series analyses within our predefined ROIs showed no significant temporal clusters differentiating the ADHD group from the TD controls (Fig. 2C). Subsequent whole-scalp cluster analyses confirmed the absence of any significant spatial group differences (Fig. 2D). This dissociation indicates that the demand-related attentional deficits observed in ADHD are driven specifically by failures in oscillatory alpha modulation, rather than by generalized shifts in aperiodic background dynamics.

### Aberrant distractor-evoked aperiodic modulation in ADHD

Building on our finding that children with ADHD exhibit deficient demand-related alpha modulation, we next sought to investigate whether they also show altered distractor-related neural responses to task-irrelevant sounds. We isolated this reactive neural response by subtracting the metrics of standard sound trials from those of deviant sound trials (Deviant minus Standard) under both low and high visual demands.

Again, we first examined oscillatory alpha power in response to auditory distractors (Fig. 3A). In contrast to the demand-related effects, time-series analyses of the alpha band (Deviant – Standard) revealed no significant reactive modulation differences between the ADHD and TD groups in both conditions. Specifically, neither group exhibited a differential oscillatory alpha response to deviant versus standard tones, with no significant interaction clusters observed in either the frontal or posterior ROIs. Exploratory whole-scalp cluster-based permutation analyses of alpha power modulation further revealed the absence of any significant spatial clusters differentiating the two groups across the scalp (Fig. 3B). This suggests that the distractor processing is not primarily driven by the oscillatory alpha system.

**Fig. 3.**
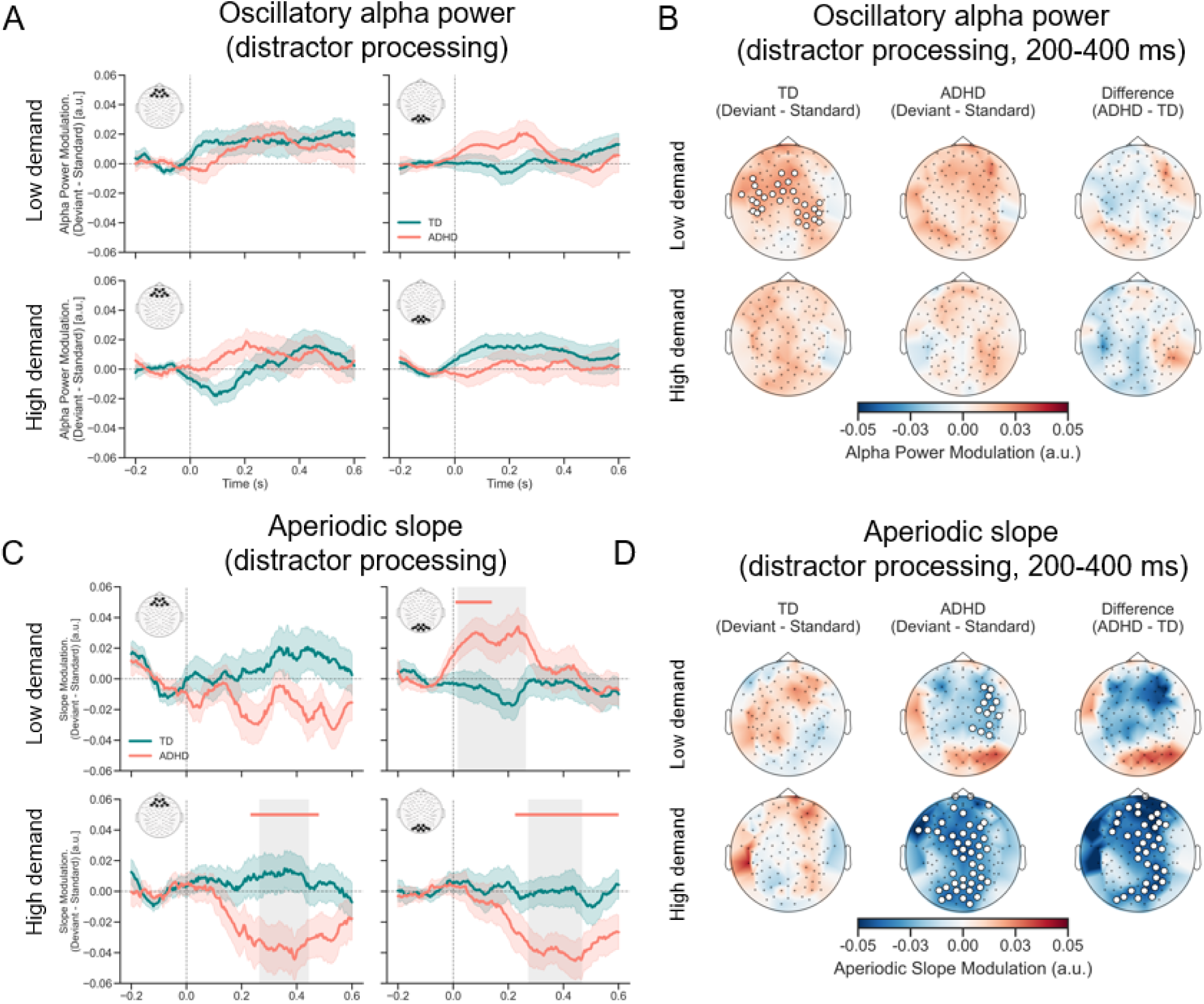
Aberrant distractor-evoked aperiodic modulation in children with ADHD and its behavioural relevance. (A) Alpha power modulation to auditory distractors (Time courses). Time courses of oscillatory alpha power modulation (Deviant minus Standard sound) under low and high visual task demands within the a priori ROIs. Shaded error bands represent the standard error (SE). (B) Alpha power modulation to auditory distractors (Topographical maps). Spatial distribution of oscillatory alpha power modulation (Deviant minus Standard sound) across the scalp for both groups, averaged over the 200–400 ms. (C) Aperiodic slope modulation to auditory distractors (Time courses). Time courses of distractor-evoked aperiodic slope modulation (Deviant minus Standard sound). Solid coloured horizontal lines indicate significant within-group temporal clusters, and grey shaded areas denote significant time windows for the group interaction. (D) Aperiodic slope modulation to auditory distractors (Topographical maps). Spatial distribution of aperiodic slope modulation and the statistical group difference averaged over the 200–400 ms. White dots denote significant spatial clusters.

Instead, the neural reactivity to auditory distractors was robustly captured by shifts in the aperiodic slope, potentially reflecting transient fluctuations in the E/I balance. In TD children, the aperiodic slope modulation (Deviant – Standard) remained negligible (near zero) across both demand conditions in the frontal and posterior ROIs (Fig. 3C, teal lines), indicating a resilient neural background (i.e., a stable cortical E/I balance) that remained unperturbed by task-irrelevant auditory changes.

Children with ADHD, however, exhibited a highly atypical and context-dependent pattern of distraction-related aperiodic modulation. Specifically, under low visual attention demand (Fig. 3C, top row), time-series analyses revealed that the ADHD group (orange lines) demonstrated a transient positive slope modulation (Deviant > Standard) within the posterior ROI (top-right panel), emerging immediately after stimulus onset and lasting until approximately 220 ms. Group comparison showed a significant difference between the two groups from 16 to 264 ms. In contrast, the frontal ROI (top-left panel) remained unaffected. Exploratory whole-scalp cluster testing, however, did not reveal a corresponding widespread posterior cluster in children with ADHD, although a small significant cluster was observed over right central scalp regions (Fig. 3D, top-middle panel). Furthermore, the direct group comparison (ADHD vs. TD) did not yield any significant spatial clusters at the whole-scalp level, suggesting that this aberrant early reactivity in the low-demand context in children with ADHD was relatively weak and spatially restricted.

Crucially, when the visual task demand increased, the reactive neural state of children with ADHD shifted drastically. Under high visual attention demand (Fig. 3C, bottom row), the ADHD group flipped their response pattern, exhibiting a sustained negative slope modulation (Deviant < Standard) in response to auditory distractor changes. Direct group comparison revealed significant differences between the two groups in both frontal (264 to 444 ms; bottom-left panel) and posterior (272 to 468 ms; bottom-right panel) ROI, indicating an atypical reactive modulation to task-irrelevant sounds. In contrast to low demand condition, topographical mapping confirmed this massive spatial divergence, revealing widespread significant negative clusters across nearly the entire scalp in the ADHD group compared to TD controls (Fig. 3D, bottom-right panel). This demand-dependent fluctuation in aperiodic activity suggests that children with ADHD may suffer from an unstable neural E/I balance, leading to dysregulation of cortical excitability when forced to handle sudden auditory distractors alongside demanding visual tasks.

### Machine learning reveals discriminative value of oscillatory and aperiodic signatures

To examine whether the identified neural abnormalities hold discriminative value for ADHD, we employed a multivariate machine learning approach using a linear support vector machine (SVM) classifier. Based on the pattern of our univariate findings, we defined three neural feature sets: sensitive neural, control neural, and all neural features (Table 1). The sensitive neural set comprised six features derived from the neural signatures highlighted by the preceding analyses: two demand-related alpha modulation features (frontal and posterior ROIs) and four distractor-related aperiodic slope features (two ROIs × two demand conditions). The control neural set comprised six separate features derived from the alternative neural measures: two demand-related aperiodic slope features and four distractor-related alpha modulation features. Finally, the all-neural set combined the sensitive and control feature sets, yielding 12 features in total (Table 1).

**Table 1.** Neural features included in the SVM classification models.

| Feature set | Feature |
|---|---|
| Sensitive neural (6) | Demand-related alpha modulation, frontal ROI |
|  | Demand-related alpha modulation, posterior ROI |
|  | Distractor-related aperiodic slope, frontal ROI, low demand |
|  | Distractor-related aperiodic slope, posterior ROI, low demand |
|  | Distractor-related aperiodic slope, frontal ROI, high demand |
|  | Distractor-related aperiodic slope, posterior ROI, high demand |
| Control neural (6) | Demand-related aperiodic slope, frontal ROI |
|  | Demand-related aperiodic slope, posterior ROI |
|  | Distractor-related alpha modulation, frontal ROI, low demand |
|  | Distractor-related alpha modulation, posterior ROI, low demand |
|  | Distractor-related alpha modulation, frontal ROI, high demand |
|  | Distractor-related alpha modulation, posterior ROI, high demand |
| All neural (12) | All 12 features listed above |

Using a repeated stratified 3-fold cross-validation procedure, we evaluated the ability of these feature sets to classify children with ADHD versus TD controls. As shown in Fig. 4A, the sensitive neural model achieved a mean area under the receiver operating characteristic curve (AUC) of 0.77, performing significantly above chance (FDR-corrected *p* < 0.01). In contrast, the control neural model did not perform significantly above chance (AUC = 0.57, *p* = 0.14). Additionally, we tested a combined model including all 12 neural features and found that its AUC (0.74, FDR-corrected *p* < 0.01) was numerically lower than sensitive model alone, suggesting that integrating non-sensitive features did not help classification performance. Statistical comparisons confirmed that the sensitive model significantly outperformed the control model (FDR-corrected *p* < 0.01). Furthermore, we examined classification performance based on task behavioral outcomes (RT and d′) alone and in combination with the neural feature sets (Fig. 4B). Behavioral features alone showed significant classification performance (AUC = 0.73, FDR-corrected *p* < 0.01). Adding the control neural features numerically reduced the AUC to 0.70, whereas adding the sensitive neural features numerically increased it to 0.79, the highest among the models tested. The combination of behavior and sensitive neural features significantly outperformed those including all neural features (AUC = 0.75) or control neural features (AUC = 0.70), with both FDR-corrected *p*s < 0.05. These results suggest that classification performance depended on the sensitivity of the neural features included, rather than simply on adding more predictors.

**Fig. 4.**
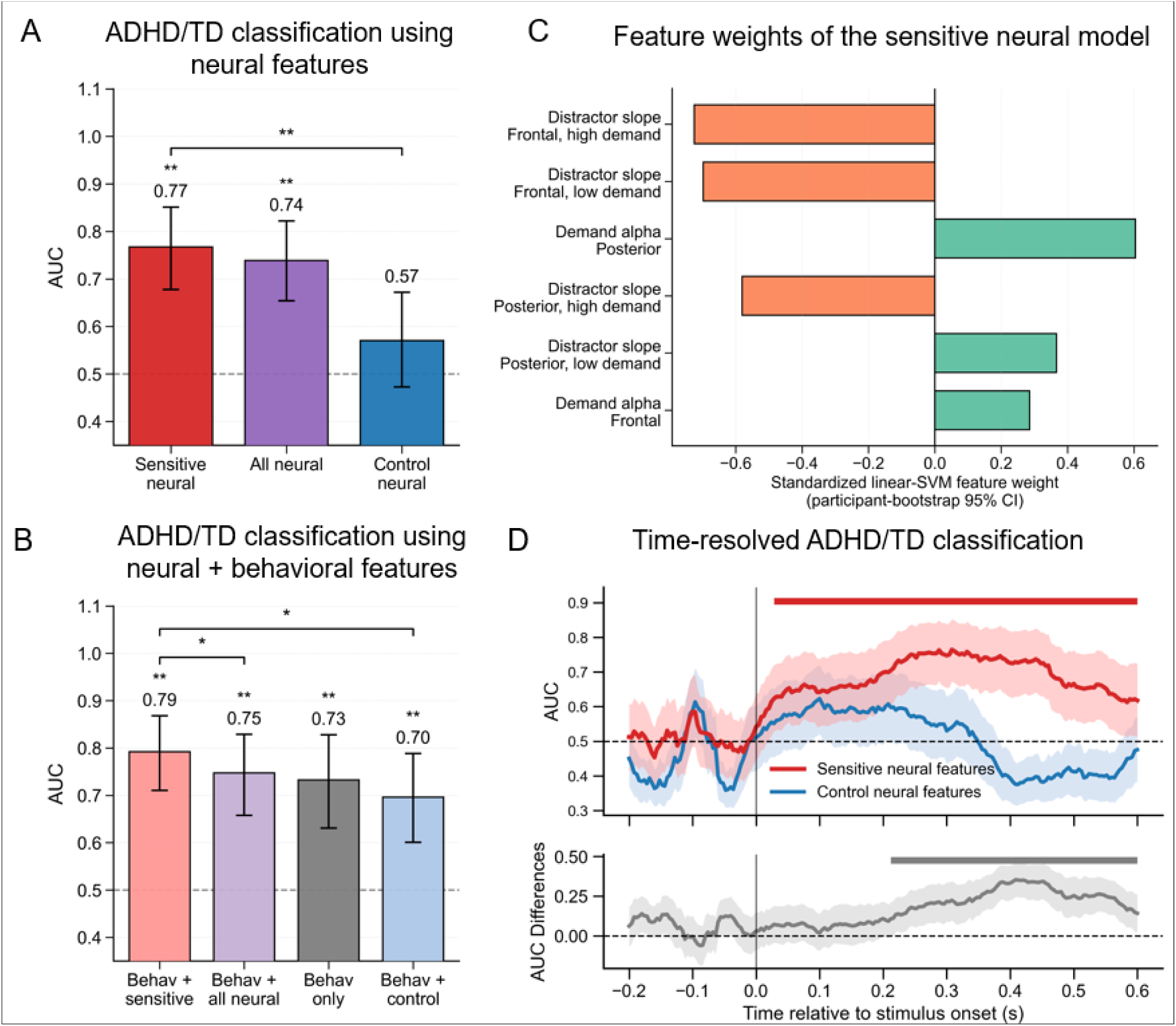
Multivariate machine learning classification of ADHD and TD groups. (A) Classification performance using neural feature sets. Comparison of the mean area under the ROC curve (AUC) among models utilizing sensitive neural features (red), all neural features (purple), and control neural features (blue). Error bars represent bootstrapped 95% confidence intervals. (B) Classification performance using behavioral and neural feature sets. Comparison of AUC scores for behavioral measures alone (reaction time and d′) and for models integrating these behavioral measures with the respective neural feature sets. (C) Feature importance for the sensitive neural model. Standardized linear-SVM weights illustrate the direction and relative contribution of each sensitive neural feature. (D) Time-resolved decoding dynamics. Top panel: Time courses of the AUC for the sensitive (red) and control (blue) neural models from –200 to 600 ms relative to stimulus onset. The solid red horizontal line indicates significant temporal clusters where the sensitive model performed significantly above chance (cluster-corrected *p* < 0.05). Bottom panel: Time course of the paired AUC difference (Sensitive minus Control). The solid gray horizontal line highlights significant temporal clusters where the sensitive model significantly outperformed the control model (cluster-corrected *p* < 0.05). Shaded error bands represent bootstrapped 95% CIs. \**p* < 0.05, \*\**p* < 0.01.

To characterize the multivariate pattern underlying the sensitive neural model (red bar in Fig. 4A), we examined the standardized linear-SVM weights of its linear-SVM decision function (Fig. 4C). In a linear SVM, the absolute magnitude of these weights provides an indication of the relative contribution of each feature to the classifier, indicating how strongly each neural signature contributes to the classifier’s ability to discriminate between the ADHD and TD groups. We found that the largest absolute weights were observed for frontal distractor-related aperiodic-slope measures and posterior demand-related alpha modulation, suggesting that these features made relatively large contributions to the multivariate classification pattern.

Furthermore, to characterize the temporal evolution of this discriminative information, we performed a time-resolved decoding analysis from –200 to 600 ms relative to stimulus onset (Fig. 4D). We found that the sensitive neural features yielded significant classification performance that emerged almost immediately after stimulus onset and sustained continuously throughout the entire processing window (cluster-corrected *p* < 0.05). Conversely, the decoding trajectory for the control neural features hovered around chance level and yielded no significant temporal clusters.

In sum, our machine learning classification analyses demonstrated that the sensitive neural features, specifically demand-related alpha and distractor-related aperiodic activity, carried greater discriminative information than the corresponding control neural features. These findings complement the univariate results by showing that identifying distinct neural mechanisms underlying task-demand adjustment and distractor reactivity could further contribute to multivariate differentiation between children with ADHD and TD controls.

## Discussion

The present study investigated how children with ADHD coordinate neural resources when facing task demand and environmental distraction. By separating oscillatory and aperiodic components of neural activity, our findings suggest that the deficit in children with ADHD during demand-related regulation is primarily captured by impaired oscillatory alpha modulation, whereas their atypical response to environmental distraction is uniquely reflected in an unstable cortical E/I balance, as shown by fluctuations in the aperiodic activity. Machine learning analyses further showed that demand-related alpha and distractor-related aperiodic activity carried greater discriminative information than the corresponding control neural features. These findings suggest that oscillatory and aperiodic neural signatures are differentially sensitive to ADHD-related alterations in distinct aspects of attentional control, and may provide potential neural markers for differentiating children with ADHD from TD controls.

Our results show that while TD children adaptively modulate alpha power to meet varying task demands, children with ADHD exhibit an atypical pattern, suggesting altered alpha-related regulation of task-relevant processing. Specifically, higher demand was associated with robust alpha desynchronization over frontal regions in TD children, consistent with previous studies linking alpha power to top-down control and task engagement [8, 43–45]. In contrast, the ADHD group not only lacked this adaptive frontal modulation, but also exhibited an unexpected increase in posterior alpha synchronization as visual demand increased. Alpha synchronization has been thought to support attentional gating by reducing the processing of currently irrelevant information (Jensen & Mazaheri, 2010; Zhao et al., 2023). In this context, posterior ERS during a demanding visual task may indicate an inappropriate inhibitory response in ADHD, potentially limiting the processing of task-relevant visual information. In addition, recent cross-modal evidence suggests that alpha modulation does not simply block early sensory input, but may help determine whether sensory information is effectively passed on to later processing stages [46]. Therefore, the abnormal posterior synchronization in children with ADHD may also reflect a broader difficulty in transmitting goal-directed information. This pattern extends previous evidence of atypical alpha modulation in ADHD by highlighting both frontal and posterior contributions to demand-related regulation [17, 18, 20]. Notably, while alpha oscillations tracked ADHD-related impairments in handling task demands, they did not differ between groups during auditory distraction processing. This dissociation suggests that alpha oscillations may be specifically sensitive to demand-related control deficits in ADHD, rather than atypical responses to environmental distraction.

Instead, the aperiodic component showed the opposite pattern: no reliable group differences were observed in its modulation by visual task demand, whereas clear differences emerged during auditory distractor processing. In TD children, the resilience of the aperiodic slope across conditions suggests a robust cortical E/I balance that remains unperturbed by irrelevant sounds [26, 33]. In contrast, the significant negative shift (steeper slope) in ADHD indicates that their neural background is more sensitive to environmental changes. This steeper slope in children with ADHD aligns with prior evidence, which suggests a compensatory increase in cortical GABAergic activity during cognitive control [47–49]. Given the excessive sensitivity to auditory distractors in ADHD, these children may fail to automatically filter out noise [3]. To manage this interruption, they may try to exert extra inhibitory resources to maintain focus, leading to a reactive increase in GABAergic signaling. This interpretation helps reconcile studies reporting a flatter slope in ADHD during No-go tasks [50]. While No-go trials require goal-directed proactive control, our task involves reactive management of environmental distraction. In No-go tasks, a flatter slope represents a primary failure to engage inhibitory control. In our study, the steeper slope may reflect an extra, over-reactive attempt to suppress a distractor that has already interfered with the task. Developmental differences may also help explain these inconsistent findings. Steeper resting-state slopes have been reported in younger ADHD samples, including children with a mean age of 5.75 years in Robertson et al. [49] and children with a mean age of 9.61 years in Dakwar-Kawar et al. [51], whereas flatter slopes were observed in adolescents with ADHD with a mean age of 13.95 years in Ostlund et al. [52]. These results suggest that ADHD cognitive impairments are linked to a deviation, with slopes that are either too flat or too steep from an optimal E/I balance at different developmental ranges [49]. In contrast, aperiodic activity did not differentiate between groups during task demand regulation, suggesting that this neural signature may be more sensitive to ADHD-related differences in the processing of task-irrelevant events.

The machine learning results further showed that these dissociable neural signatures held discriminative value for ADHD-TD differentiation at the individual level. The sensitive neural feature set, comprising six features derived from demand-related alpha modulation and distractor-related aperiodic slope, achieved higher ADHD-TD classification performance. In contrast, the matched control neural feature set, comprising demand-related aperiodic slope and distractor-related alpha modulation, showed limited discriminative performance. Importantly, adding all neural features did not further improve classification beyond the sensitive neural model; similarly, when behavioral performance was included, the broader combined-feature model showed lower classification performance than the model combining behavior with only sensitive neural features. Together, these results suggest that alpha and aperiodic activity provided the optimal discriminative information when they were aligned with the attentional processes in which ADHD-related abnormalities were most evident. The lack of additional benefit from broader feature combinations further indicates that ADHD classification may depend more on selecting features sensitive to specific pathological processes than on simply increasing the number of predictors, consistent with the view that irrelevant or redundant features may reduce model efficiency [41]. This process-specific interpretation is also consistent with recent evidence that aperiodic and alpha oscillatory activity can carry distinct neural information rather than reflecting interchangeable components of the EEG signal [25]. Notably, the SVM feature weights also revealed that a feature without a significant univariate group difference, the frontal low-demand aperiodic slope, could still contribute substantially to classification. This suggests that multivariate models may capture subtle distributed patterns that are not evident when individual features are examined separately. Thus, separating oscillatory and aperiodic dynamics across distinct attentional processes may help identify more refined, process-specific markers for ADHD differentiation.

Several limitations should be considered. While the aperiodic exponent has been established as a robust marker of the cortical E/I balance [30, 53–55], it remains an indirect measure derived from scalp EEG. Future studies combining EEG with complementary methods, such as magnetic resonance spectroscopy, may help to clarify this relationship [29, 56]. Furthermore, the current study focused on school-aged children using a specific cross-modal task, and future work will be needed to determine whether similar oscillatory and aperiodic patterns generalize across developmental stages and more naturalistic attentional settings.

In conclusion, by dissociating the oscillatory and aperiodic components of neural activity, the present study demonstrates that ADHD attention deficits may be driven by distinct neural pathways. Impairments in demand-related regulation are primarily linked to failures in alpha-mediated gating, whereas atypical responses to distraction are reflected in an unstable cortical E/I balance, as shown by aperiodic slope fluctuations. Critically, these ADHD-related neural signatures effectively supported machine-learning-based differentiation between ADHD and TD children, showing superior discriminative capacity compared to control feature sets. These findings provide a comprehensive framework for understanding ADHD as a disorder of both goal-directed gating and neural background stability, highlighting the importance of considering different dimensions of neural dynamics when investigating cognitive dysfunction and informing clinical assessment [25, 57]. Furthermore, these distinct neural signatures may offer promising targets for future clinical interventions, where targeted neuromodulation techniques could be developed to selectively regulate oscillatory or aperiodic activity [32] to improve attentional control in children with ADHD.

## Materials and Methods

### Participants

The current study utilized the same dataset from a previously reported cohort [3], but addressed distinct theoretically motivated questions concerning oscillatory and aperiodic neural activity, using analysis approaches that were not included in the original report. The final sample size was comparable to, or larger than, those reported in previous EEG studies examining alpha oscillations, aperiodic activity, and attentional processing in children with ADHD [22, 51].

We initially recruited a total of 117 children, comprising children with ADHD and TD children. ADHD diagnoses were confirmed by certified psychiatrists using the Kiddie Schedule for Affective Disorders and Schizophrenia for School-Age Children-Present and Lifetime Version (K-SADS-PL) [58], following the Diagnostic and Statistical Manual of Mental Disorders, 5th ed. (DSM-V) [59], as part of routine clinical evaluations. TD children were screened through interviews with both parents and teachers to rule out psychiatric or neurological concerns.

All participants met the following inclusion criteria: normal hearing; normal or corrected-to-normal vision; no history of neurological disease or head trauma with loss of consciousness; no major psychiatric comorbidities, including schizophrenia, severe affective disorders, autism spectrum disorders, or developmental disabilities; and a full-scale intelligence quotient (IQ) above 80, as assessed by the Wechsler Intelligence Scale for Children, 4th ed. (WISC-IV) [60]. ADHD symptom severity was quantified using parent ratings on the ADHD Rating Scale (ADHD-RS) [61]. After exclusions based on clinical criteria, excessive EEG artifacts, and trial rejection, data from 90 children were included in the final analyses (ADHD: 45, age = 10.62 ± 1.93 years, range = 7–14 years, 10 females; TD: 45, age = 10.20 ± 1.54 years, range = 7–13 years, 10 females). A detailed recruitment and exclusion flowchart is presented in Fig. S1. The two groups did not differ significantly in age, sex, handedness, or IQ, whereas children with ADHD showed significantly higher symptom severity (Table 2).

**Table 2.** Demographic information and symptom scores of the two groups of children.

|  | ADHD group | TD group | Group Difference |  |
|---|---|---|---|---|
|  | (n = 45) | (n = 45) | *χ*<sup>2</sup> or *t* | *p*-value |
| Sex (boy: girl) | 35: 10 | 35: 10 | 0.000 | 1.000 |
| Handedness (left: right) | 2: 43 | 0: 45 | 0.511 | .475 |
| Age (years) | 10.62 ± 1.93 | 10.20 ± 1.54 | 1.157 | .251 |
| Full-scale IQ | 110.09 ± 12.41 | 113.27 ± 11.60 | –1.255 | .213 |
| ADHD-RS |  |  |  |  |
| Inattention | 17.73 ± 3.09 | 7.00 ± 2.80 | 17.248<sup>***</sup> | <.001 |
| Hyperactivity/impulsivity | 11.38 ± 5.72 | 4.58 ± 2.83 | 7.149<sup>***</sup> | <.001 |

*Note.* All values are presented as the mean ± SD, unless otherwise indicated. ADHD, attention-deficit/hyperactivity disorder. TD, typically developing. ADHD-RS, subscale scores of ADHD Rating Scale. <sup>\*\*\*</sup>*p* < 0.001.

This study was approved by the Ethics Committee of Beijing Normal University, and verbal and written informed consent was obtained from all participants and their legal guardians in accordance with the Declaration of Helsinki.

### Stimuli and procedure

The experimental stimuli and procedure were identical to those described in our previous publication [3]. Children performed a visual detection task under two levels of visual attention demand (low and high), while task-irrelevant auditory oddball stimuli were presented concurrently (Fig. 1A).

Visual stimuli consisted of a circular search array including twelve items (one circle and eleven diamonds) with fixation. In 10% of the trials, the circle served as the visual target, and participants were instructed to respond as quickly and accurately as possible. Visual attention demands were manipulated by varying the perceptual similarity between targets and non-targets. In the low visual attention demand condition, the target circle was red and visually salient, whereas in the high visual attention demand condition it was dark gray and similar to the non-target color, requiring greater attentional resources to detect. No responses were required on 90% non-target trials.

Auditory stimuli were presented synchronously with the visual arrays and followed an oddball structure (200 ms duration, 600 ms inter-stimulus interval), with frequent standard tones (80%) and infrequent deviant tones (20%). Participants were instructed to ignore the auditory stimuli and focus on the visual task. The proportion of standard and deviant tones was identical across visual target and non-target trials. Each of the two conditions contained eight blocks of 100 trials, which were performed separately, with the order of conditions counterbalanced between participants.

### EEG recording and data processing

Continuous EEG signals were recorded using a 128-channel EGI system (Electrical Geodesics, Inc.) at a sampling rate of 1000 Hz. During acquisition, the signals were amplified using an online 0.01– 400 Hz bandpass filter. The Cz electrode served as the online reference, and electrode impedances were maintained below 50 kΩ.

Data pre-processing was performed in MATLAB using the EEGLAB toolbox [62]. Initially, the 20 outermost electrodes susceptible to movement artifacts were excluded prior to analysis, and the remaining data were downsampled to 250 Hz and re-referenced to the average of the mastoids. A bandpass filter (0.5–30 Hz) was applied to the continuous data. Following the manual excision of artifact-contaminated segments and interpolation of noisy channels, independent component analysis (ICA) was subsequently employed to identify and remove ocular artifacts, including blinks and saccades.

Following ICA, the continuous EEG data were segmented into epochs spanning –600 to 800 ms relative to the stimulus onset. To guarantee robust data quality and further eliminate residual muscular or electrooculographic (EOG) interference, we applied an amplitude threshold, automatically discarding any epochs where voltage fluctuations exceeded ±100 μV at any electrode. Furthermore, to avoid confounding motor execution effects, we restricted all subsequent analyses exclusively to clean epochs from the 90% of trials that required no button response. Participants who retained fewer than 50% of their trials after artifact rejection were excluded from the final sample, resulting in the removal of six children with ADHD and three TD children.

### Oscillatory and aperiodic activity analysis

To isolate induced (non-phase-locked) neural activity from evoked responses, we first subtracted condition-specific event-related potentials (ERPs) from each trial for each participant. This procedure was intended to minimize the contribution of evoked responses to subsequent spectral parameterization. We then decomposed the ERP-subtracted data into time-frequency representations using complex Morlet wavelets. We extracted power estimates across –600 to 800 ms relative to stimulus onset, analyzing frequencies from 1 to 30 Hz in 1–Hz increments. To optimize the balance between temporal and spectral resolution, we dynamically scaled the number of wavelet cycles to half of the respective frequency (*f*/2). Following this decomposition, we averaged the power estimates across trials within each condition for every participant.

To disentangle true oscillatory neural activity from broadband aperiodic background dynamics, we applied the time-resolved spectral parameterization (FOOOF) algorithm [26] to the trial-averaged power spectra at each time point and electrode. We configured the fitting algorithm with a fixed aperiodic mode, a peak width limit of 2 to 6 Hz, a maximum of 4 peaks, and a peak threshold of 2.0. By fitting these models across the 1 to 30 Hz frequency range, we parameterized the power spectra into distinct aperiodic and periodic components. From the aperiodic fit, we extracted the aperiodic exponent and defined the aperiodic slope as the negative exponent. The periodic component, representing true neural oscillations, was obtained by subtracting the aperiodic fit from the original power spectrum.

In line with our primary hypotheses, we focused our subsequent analyses on two specific neural metrics: oscillatory alpha power (8–12 Hz) and the aperiodic slope. We baseline-corrected the time-series data for both metrics using a pre-stimulus window of –200 to 0 ms. To evaluate the spatio-temporal dynamics of these target metrics, we defined two *a priori* spatial regions of interest (ROIs) based on previous literature: a frontal ROI (electrodes E4, E5, E10, E11, E12, E18, E19, E24, E124) and a posterior ROI (electrodes E69, E70, E71, E74, E75, E76, E82, E83, E89). To assess proactive visual attention, we calculated the difference in baseline-corrected neural metrics between the high and low visual demand conditions (High minus Low) across all trials. Conversely, to evaluate reactive processing of task-irrelevant distractors, we computed the difference between trials with deviant and standard tones (Deviant minus Standard) under varying visual demands. For topographical visualizations, we averaged the neural responses over a predefined post-stimulus target window of 200 to 400 ms, as this period is associated with the active deployment of top-down attention and distractor suppression [42, 63].

### Multivariate machine learning classification

To determine the discriminative value of the identified distinct neural signatures, we trained linear SVM classifiers to distinguish children with ADHD from TD controls. Building upon our univariate findings, we defined two primary neural feature sets. The *sensitive neural features* set included neural metrics that exhibited significant group differences: demand-related alpha power modulation (high minus low demand) and distractor-related aperiodic slope modulation (deviant minus standard sound) extracted from the frontal and posterior ROIs, yielding 6 features in total (2 demand-related alpha features and 4 distractor-related slope features across two demand conditions). For comparison, we constructed a *control neural features* set containing 6 features that showed no significant group differences in the univariate analyses: demand-related aperiodic slope modulation and distractor-related alpha power modulation from the same ROIs.

For the static classification, neural features were derived from the predefined 200–400 ms time window. To comprehensively evaluate classification performance, we also integrated behavioral features (average visual task discriminability d′ and reaction time) to construct models with combined feature sets. The classification pipeline was implemented using *scikit-learn* in Python. To prevent data leakage, feature standardization (z-scoring) was nested within the cross-validation loop, such that the scaler was fitted solely on the training folds and subsequently applied to the corresponding test folds. The linear SVM classifiers were configured with balanced class weights to account for any slight sample size imbalances within individual training folds. Model performance was evaluated using a repeated stratified 3-fold cross-validation scheme (100 repeats). The primary evaluation metric was the mean area under the receiver operating characteristic curve (AUC), calculated separately within each test fold and then averaged across all repeats and folds to prevent information leakage and ensure unbiased estimates.

To characterize the temporal dynamics of discriminative information, we extended our classification approach to a time-resolved decoding analysis spanning from –200 to 600 ms relative to stimulus onset. At each sampling time point, independent linear SVM classifiers were trained and tested using the same repeated stratified 3-fold cross-validation procedure.

### Statistical analysis

To address the multiple comparisons problem inherent to high-dimensional EEG data, we evaluated statistical significance of univariate neural responses using non-parametric cluster-based permutation testing. For both topographical maps and ROI time series, this approach allowed us to identify significant clusters of neural activity and decoding performance without *a priori* assumption regarding exact spatial or temporal bounds. We assessed within-group effects, such as the neural modulation to varying visual demands (High minus Low) and the reactivity to irrelevant sounds (Deviant minus Standard), using one-sample cluster permutation tests against zero (two-tailed). For all permutation tests, we generated null distributions using 1000 random permutations and defined significant clusters using an alpha threshold of 0.05. Spatial clustering was defined according to a sensor neighborhood graph, while temporal clustering was applied continuously across the predefined post-stimulus epochs. For the multivariate machine learning classification, we employed non-parametric resampling techniques to assess both statistical significance and model uncertainty. First, to test whether each static model’s mean AUC was significantly higher than chance level (0.5), we conducted full-pipeline label permutation tests (1000 iterations), where diagnostic labels were randomly shuffled prior to the cross-validation loop. Second, to construct 95% confidence intervals for both the individual model AUCs and the paired AUC differences, we performed a stratified participant-level bootstrap procedure (1000 iterations) on the obtained cross-fitted scores. Third, for planned pairwise model comparisons, we implemented participant-wise paired score-swap permutation tests (1000 iterations). To place competing models on a common scale before score swapping, each participant’s cross-fitted decision scores were first transformed into fold-normalized percentile ranks; their entire repeat-wise score trajectories were then randomly swapped between the two models to generate the null distribution of AUC differences under the one-tailed alternative. Multiple comparisons across the family of planned model evaluations and chance-level tests were controlled using the FDR correction.

For the time-resolved decoding analysis, we applied cluster-based permutation tests to control for multiple comparisons across the temporal dimension. Pointwise 95% CIs for both the time-resolved AUC curves and their difference were estimated using a stratified participant-level bootstrap (1000 iterations). To evaluate whether decoding performance was significantly above chance over time, null distributions of AUC curves were generated using full-pipeline label permutations (1000 iterations). Similarly, to test the temporal intervals where the sensitive model outperformed the control model (Sensitive > Control), null distributions of AUC difference curves were generated using participant-wise score-swap permutations (1000 iterations) of the fold-normalized percentile ranks.

### Ethical Approval

This study was approved by the Ethics Committee of Beijing Normal University, and verbal and written informed consent was obtained from all participants and their legal guardians in accordance with the Declaration of Helsinki.

## Supplemental Materials

**Figure S1.**
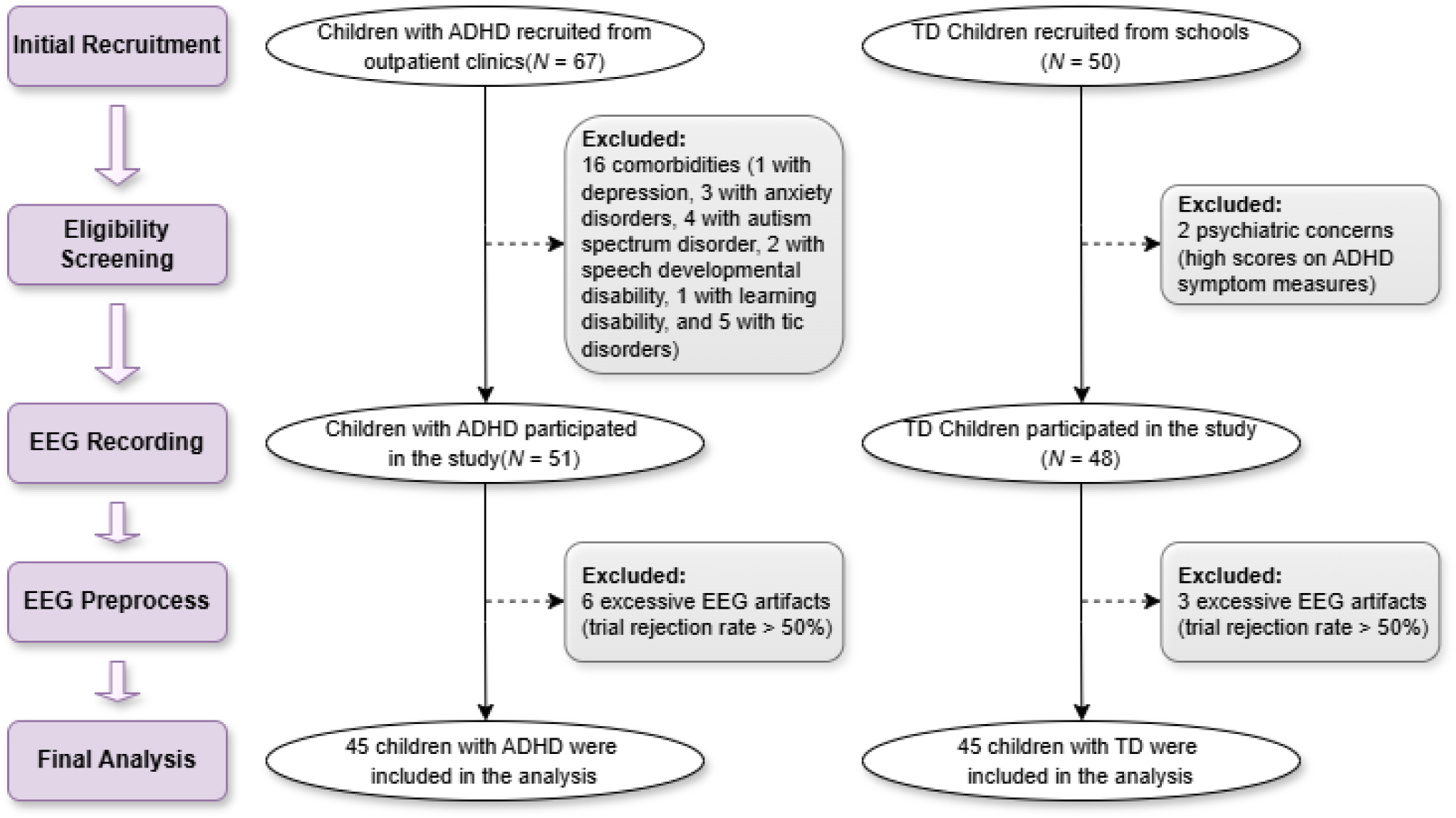
Flowchart of participant inclusion and exclusion. ADHD = attention-deficit/hyperactivity disorder; EEG = electroencephalography; TD = typically developing.

## References

1. Thomas R, Sanders S, Doust J, Beller E, Glasziou P. Prevalence of attention-deficit/hyperactivity disorder: a systematic review and meta-analysis. Pediatrics. 2015;135(4):e994–e1001.

2. Klein JL, Allen HA, Clibbens J, Cook A, Amanatidou V, Mavritsaki E. Selective visual attention in ADHD: a narrative review. Curr Neurol Neurosci Rep. 2025;25(1):51.

3. Kong Y, Yuan X, Sun L, et al. Altered processing of auditory distractions under competing inputs in children with attention-deficit/hyperactivity disorder. J Am Acad Child Adolesc Psychiatry. Published online November 13, 2025.

4. Friedman-Hill SR, Wagman MR, Gex SE, Pine DS, Leibenluft E, Ungerleider LG. What does distractibility in ADHD reveal about mechanisms for top-down attentional control? Cognition. 2010;115(1):93–103.

5. Rivella C, Bazzurro A, Olla V, et al. Impact of distractors on sustained attention and inhibition in children with ADHD. J Atten Disord. 2024;28(6):970–981.

6. Wang E, Sun L, Sun M, et al. Attentional selection and suppression in children with attention-deficit/hyperactivity disorder. Biological Psychiatry: Cognitive Neuroscience and Neuroimaging. 2016;1(4):372–380.

7. Jensen O, Mazaheri A. Shaping functional architecture by oscillatory alpha activity: gating by inhibition. Front Hum Neurosci. 2010;4.

8. Klimesch W, Sauseng P, Hanslmayr S. EEG alpha oscillations: the inhibition–timing hypothesis. Brain Research Reviews. 2007;53(1):63–88.

9. Zhao C, Kong Y, Li D, et al. Suppression of distracting inputs by visual-spatial cues is driven by anticipatory alpha activity. PLOS Biology. 2023;21(3):e3002014.

10. Bauer M, Stenner MP, Friston KJ, Dolan RJ. Attentional modulation of alpha/beta and gamma oscillations reflect functionally distinct processes. J Neurosci. 2014;34(48):16117–16125.

11. Klimesch W. Alpha-band oscillations, attention, and controlled access to stored information. Trends Cogn Sci. 2012;16(12):606–617.

12. Zhu Y, Wang Q, Zhang L. Study of EEG characteristics while solving scientific problems with different mental effort. Sci Rep. 2021;11(1):23783.

13. Mazaheri A, van Schouwenburg MR, Dimitrijevic A, Denys D, Cools R, Jensen O. Region-specific modulations in oscillatory alpha activity serve to facilitate processing in the visual and auditory modalities. NeuroImage. 2014;87:356–362.

14. van Diepen RM, Mazaheri A. Cross-sensory modulation of alpha oscillatory activity: suppression, idling, and default resource allocation. European Journal of Neuroscience. 2017;45(11):1431–1438.

15. Foxe JJ, Simpson GV, Ahlfors SP. Parieto-occipital ∼10Hz activity reflects anticipatory state of visual attention mechanisms. NeuroReport. 1998;9(17):3929.

16. Fu KMG, Foxe JJ, Murray MM, Higgins BA, Javitt DC, Schroeder CE. Attention-dependent suppression of distracter visual input can be cross-modally cued as indexed by anticipatory parieto–occipital alpha-band oscillations. Cognitive Brain Research. 2001;12(1):145–152.

17. Lenartowicz A, Mazaheri A, Jensen O, Loo SK. Aberrant modulation of brain oscillatory activity and attentional impairment in attention-deficit/hyperactivity disorder. Biological Psychiatry: Cognitive Neuroscience and Neuroimaging. 2018;3(1):19–29.

18. Mazaheri A, Fassbender C, Coffey-Corina S, Hartanto TA, Schweitzer JB, Mangun GR. Differential oscillatory electroencephalogram between attention-deficit/hyperactivity disorder subtypes and typically developing adolescents. Biological Psychiatry. 2014;76(5):422–429.

19. Michelini G, Salmastyan G, Vera JD, Lenartowicz A. Event-related brain oscillations in attention-deficit/hyperactivity disorder (ADHD): A systematic review and meta-analysis. International Journal of Psychophysiology. 2022;174:29–42.

20. Arjona A, Angulo-Ruiz BY, Rodríguez-Martínez EI, Cabello-Navarro C, Gómez CM. Time-frequency neural dynamics of ADHD children and adolescents during a working memory task. Neuroscience Letters. 2023;798:137100.

21. 21. Huurne N ter, Onnink M, Kan C, Franke B, Buitelaar J, Jensen O. Behavioral consequences of aberrant alpha lateralization in attention-deficit/hyperactivity disorder. Biological Psychiatry. 2013;74(3):227–233.

22. Vollebregt MA, Zumer JM, ter Huurne N, Buitelaar JK, Jensen O. Posterior alpha oscillations reflect attentional problems in boys with attention deficit hyperactivity disorder. Clin Neurophysiol. 2016;127(5):2182–2191.

23. Cañigueral R, Palmer J, Ashwood KL, et al. Alpha oscillatory activity during attentional control in children with autism spectrum disorder (ASD), attention-deficit/hyperactivity disorder (ADHD), and ASD+ADHD. Journal of Child Psychology and Psychiatry. 2022;63(7):745–761.

24. Zhao J, Bao M, Ruan W, et al. Electrophysiological abnormalities associated with sustained attention in children with attention deficit hyperactivity disorder and autism spectrum disorder. IEEE Transactions on Neural Systems and Rehabilitation Engineering. 2025;33:1785–1795.

25. Bender A, Zhao C, Vogel E, Awh E, Voytek B. Differential representations of spatial location by aperiodic and alpha oscillatory activity in working memory. Proc Natl Acad Sci USA. 2025;122(30):e2506418122.

26. Donoghue T, Haller M, Peterson EJ, et al. Parameterizing neural power spectra into periodic and aperiodic components. Nat Neurosci. 2020;23(12):1655–1665.

27. Lu R, Dermody N, Duncan J, Woolgar A. Aperiodic and oscillatory systems underpinning human domain-general cognition. Commun Biol. 2024;7(1):1643.

28. Lu R, Pollitt E, Woolgar A. Distinct and complementary mechanisms of oscillatory and aperiodic alpha activity in visuospatial attention. Imaging Neuroscience. 2025;3:IMAG.a.1038.

29. Cohen Kadosh R. Rethinking excitation/inhibition balance in the human brain. Nat Rev Neurosci. 2025;26(8):451–452.

30. Gao R, Peterson EJ, Voytek B. Inferring synaptic excitation/inhibition balance from field potentials. NeuroImage. 2017;158:70–78.

31. Lendner JD, Helfrich RF, Mander BA, et al. An electrophysiological marker of arousal level in humans. Elife. 2020;9:e55092.

32. Lu R, Rogan IW, Woolgar A. Optimizing control conditions for entraining neural oscillations using rhythmic TMS. Brain Stimulation. 2025;18(5):1367–1369.

33. Voytek B, Kramer MA, Case J, et al. Age-related changes in 1/ *f* neural electrophysiological noise. J Neurosci. 2015;35(38):13257–13265.

34. Gyurkovics M, Clements GM, Low KA, Fabiani M, Gratton G. Stimulus-induced changes in 1/f-like background activity in EEG. J Neurosci. 2022;42(37):7144–7151.

35. Kałamała P, Gyurkovics M, Bowie DC, et al. Event-induced modulation of aperiodic background EEG: attention-dependent and age-related shifts in E:I balance, and their consequences for behavior. Imaging Neuroscience. 2024;2:imag-2-00054.

36. Donoghue T. A systematic review of aperiodic neural activity in clinical investigations. European Journal of Neuroscience. 2025;62(7):e70255.

37. Sonuga-Barke EJS, Wiersema JR, van der Meere JJ, Roeyers H. Context-dependent dynamic processes in attention deficit/hyperactivity disorder: differentiating common and unique rffects of state regulation deficits and delay aversion. Neuropsychol Rev. 2010;20(1):86–102.

38. Valo S, Tannock R. Diagnostic Instability of DSM–IV ADHD Subtypes: Effects of Informant Source, Instrumentation, and Methods for Combining Symptom Reports. Journal of Clinical Child & Adolescent Psychology. 2010;39(6):749–760.

39. Wolraich ML, Hagan JF Jr, Allan C, et al. Clinical Practice Guideline for the Diagnosis, Evaluation, and Treatment of Attention-Deficit/Hyperactivity Disorder in Children and Adolescents. Pediatrics. 2019;144(4):e20192528.

40. Cao M, Martin E, Li X. Machine learning in attention-deficit/hyperactivity disorder: new approaches toward understanding the neural mechanisms. Transl Psychiatry. 2023;13(1):236.

41. Pudjihartono N, Fadason T, Kempa-Liehr AW, O’Sullivan JM. A Review of Feature Selection Methods for Machine Learning-Based Disease Risk Prediction. Front Bioinform. 2022;2.

42. Zheng Y, Lu R, Woolgar A. Radical flexibility of neural representation in frontoparietal cortex and the challenge of linking it to behaviour. Current Opinion in Behavioral Sciences. 2024;57:101392.

43. Bonnefond M, Jensen O. The role of alpha oscillations in resisting distraction. Trends in Cognitive Sciences. 2025;29(4):368–379.

44. Foxe JJ, Snyder AC. The role of alpha-band brain oscillations as a sensory suppression mechanism during selective attention. Front Psychol. 2011;2.

45. Jensen O, Bonnefond M. The alpha rhythm: from physiology to behavior. Physiol Rev. 2026;106(3):1123–1159.

46. 46. Brickwedde M, Limachya R, Markiewicz R, et al. Cross-modal interaction of human alpha activity does not reflect inhibition of early sensory processing in a frequency-tagging study using EEG and MEG. Groen II, Luo H, eds. eLife. 2026;14:RP106050.

47. Pani SM, Saba L, Fraschini M. Clinical applications of EEG power spectra aperiodic component analysis: a mini-review. Clinical Neurophysiology. 2022;143:1–13.

48. Peterson EJ, Rosen BQ, Belger A, Voytek B, Campbell AM. Aperiodic neural activity is a better predictor of schizophrenia than neural oscillations. Clin EEG Neurosci. 2023;54(4):434–445.

49. Robertson MM, Furlong S, Voytek B, Donoghue T, Boettiger CA, Sheridan MA. EEG power spectral slope differs by ADHD status and stimulant medication exposure in early childhood. Journal of Neurophysiology. 2019;122(6):2427–2437.

50. Pertermann M, Bluschke A, Roessner V, Beste C. The modulation of neural noise underlies the effectiveness of methylphenidate treatment in attention-deficit/hyperactivity disorder. Biological Psychiatry: Cognitive Neuroscience and Neuroimaging. 2019;4(8):743–750.

51. Dakwar-Kawar O, Mentch-Lifshits T, Hochman S, et al. Aperiodic and periodic components of oscillatory brain activity in relation to cognition and symptoms in pediatric ADHD. Cereb Cortex. 2024;34(6):bhae236.

52. Ostlund BD, Alperin BR, Drew T, Karalunas SL. Behavioral and cognitive correlates of the aperiodic (1/*f*-like) exponent of the EEG power spectrum in adolescents with and without ADHD. Developmental Cognitive Neuroscience. 2021;48:100931.

53. Chaudhuri R, He BJ, Wang XJ. Random recurrent networks near criticality capture the broadband power distribution of human ECoG dynamics. Cereb Cortex. 2018;28(10):3610–3622.

54. Lu R. Linking the multiple-demand cognitive control system to human electrophysiological activity. Neuropsychologia. 2025;210:109096.

55. Trakoshis S, Martínez-Cañada P, Rocchi F, et al. Intrinsic excitation-inhibition imbalance affects medial prefrontal cortex differently in autistic men versus women. eLife. 2020;9:e55684.

56. McKeon SD, Perica MI, Parr AC, et al. Aperiodic EEG and 7T MRSI evidence for maturation of E/I balance supporting the development of working memory through adolescence. Developmental Cognitive Neuroscience. 2024;66:101373.

57. Van Engen Q, Chau G, Smith A, Adam K, Donoghue T, Voytek B. Dissociating contributions of theta and alpha oscillations from aperiodic neural activity in human visual working memory. J Neurosci. 2026;46(7):e2340242025.

58. Kaufman J, Birmaher B, Brent D, et al. Schedule for affective disorders and schizophrenia for school-age children-present and lifetimeversion (K-SADS-PL): initial reliability and validity data. Journal of the American Academy of Child & Adolescent Psychiatry. 1997;36(7):980–988.

59. American Psychiatric Association. Diagnostic and Statistical Manual of Mental Disorders. 5th ed. Arlington: American Psychiatric Publishing, Inc.; 2013.

60. Wechsler D. Wechsler Intelligence Scale for Children. 4th ed. New York: The Psychological Corporation; 2003.

61. DuPaul GJ, Power TJ, Anastopoulos AD, Reid R. ADHD Rating Scale—IV: Checklists, Norms, and Clinical Interpretation. 1st ed. New York: The Guilford Press; 1998.

62. Delorme A, Makeig S. EEGLAB: an open source toolbox for analysis of single-trial EEG dynamics including independent component analysis. Journal of Neuroscience Methods. 2004;134(1):9–21.

63. Van Moorselaar D, Slagter HA. Inhibition in selective attention. Annals of the New York Academy of Sciences. 2020;1464(1):204–221.

